# Transgenerational inheritance of betaine-induced epigenetic alterations in estrogen-responsive IGF-2/IGFBP2 genes in rat hippocampus

**DOI:** 10.1101/640524

**Authors:** Yang Yang, Shu Yang, Wenduo Jiang, Fulei Qi, Ruqian Zhao

**Author notes:** Address proofs and correspondence to: Dr. Ruqian Zhao, Key Laboratory of Animal Physiology & Biochemistry, Nanjing Agricultural University, Nanjing 210095, P. R. China.

## Abstract

Betaine as a methyl donor participates in DNA methylation. Here, we sought to investigate the effects of betaine on hippocampal expression of neurogenesis genes and their DNA methylation status across three generations. Pregnant rats (F0) were fed control and betaine-supplemented diets throughout gestation and lactation; and female F1 and F2 offspring at weaning, together with the F0 dams, were used in the study. Hippocampal expression of aromatase, estrogen receptor α and estrogen related receptor β was down-regulated in F1, together with the estrogen-responsive IGF-2/IGFBP2 genes. However, all these genes were up-regulated in F2, which follows the same pattern of F0. In agreement with changes in mRNA expression, the imprinting control region (ICR) of IGF-2 gene was hypomethylated in F1 but hypermethylated in F2 and F0. In contrast, the promoter DNA methylation status of all the affected genes was hypermethylated in F1 but hypomethylated in F2 and F0. Betaine homocysteine methyltransferase followed same pattern of transgenerational inheritance. These results indicate that betaine exerts a transgenerational effect on hippocampal expression of estrogen-responsive genes in rat offspring, which is associated with corresponding alterations in DNA methylation on ICR of IGF-2 gene and the promoter of affected genes.

**Author Summary:** Epigenetic transgenerational inheritance is defined as the phenotypic change across generations and germline transmission of epigenetic information without direct genetic manipulation or environmental exposure. Maternal and paternal diet has been shown to have transgenerational behavioral and physiological effects. Maternal intake of methyl donors, including folate, betaine, and choline during different gestation period alters epigenetic modifications of many imprinted genes in the genomic DNA isolated from cord blood. Betaine as a methyl donor nutrition plays an important role in brain development. However, the effect of maternal betaine supplementation on transgenerational inheritance of imprinted gene IGF-2 in offspring hippocampus is still unknown. Here, we sought to investigate the effects of betaine on rat hippocampal expression of neurogenesis genes and their DNA methylation status across three generations. We found that betaine significantly decreased estrogen-responsive genes IGF-2/IGFBP2 expression in rat F1 offspring, yet increased in F2 offspring, which is associated with corresponding alterations in DNA methylation on ICR of IGF-2 gene and the promoter of affected genes. Our results suggesting that maternal methyl donor nutrition supplementation may be exert an epigenetic transgenerational effect on female offspring.

## 1. Introduction

Hippocampus is the key structure governing many important functions including learning and memory, as well as navigation and spatial orientation (1). Accumulated studies have shown that estrogen enhances synaptic plasticity, such as enhances long-term potentiation (LTP), the electrophysiological parameter of memory, and improves performance on hippocampus-dependent cognitive behavior (2, 3). Growing evidences indicate that 17β-estradiol can be de novo synthesized in the hippocampus (4, 5), and hippocampus-derived estradiol has been shown to be essential for synaptic plasticity (4, 6). Aromatase (CYP19A1), the final enzyme of estradiol biosynthesis, is a key mammalian microsomal enzyme in the process of the conversion of androgens to estrogens (7). Numerous studies have shown that estradiol upregulates synaptic proteins in the hippocampal CA1 region (8, 9). Vice versa, inhibition of aromatase activity significantly reduced the expression of synaptic proteins, which was associated with synapse loss in hippocampal slice cultures and in female mice (10). Moreover, the beneficial effects of estrogens on memory are mediated through classical estrogen receptors (ERs), such as ERα and ERβ, which are expressed throughout the brain (11).

It was reported that insulin-like growth factor 2 (IGF-2) and insulin-like growth factor binding protein 2 (IGFBP2) as estrogen-response gene in rat hippocampus (12). IGF-2 is an important component of the insulin/IGF-signaling axis, which plays a critical role in hippocampal neurogenesis and dendritic plasticity (13). IGF receptors are expressed throughout the CNS. Especially, IGF-2 receptor is mainly present in the hippocampus (14). IGF binding proteins were thought to modulate the biological actions of the IGF. In particular, IGFBP2 is assumed to play a dominant role in the regulation of IGF in the CNS which is the most abundant IGFBP expressed in the brain (15, 16). Additionally, IGFBP2 was reported that contains two heparin-binding domains (HBD) can bind IGF-2 to form IGF-2/IGFBP2 complex to protecting IGF-2 from clearance by IGF2R mediated endocytosis (17).

IGF-2 and H19 are imprinted genes exhibiting reciprocity in allele-specific expression (18, 19). Monoallelic expression of the two genes is regulated by a differentially methylated region (DMR) also named as imprinting control region (ICR), are differentially methylated between paternal and maternal alleles (20). The IGF2/H19 gene cluster is regulated by an ICR that contains multiple binding sites for the insulator protein, CCCTC binding factor (CTCF) (21). On the maternal allele, an unmethylated IGF2/H19 ICR binding of CTCF prevents the transcription of IGF-2 and activates H19. On the paternal allele, highly methylated IGF2/H19 ICR binding of CTCF activates transcription of IGF2 and silences H19. Previous studies reported that maternal intake of methyl donors, including folate, betaine, and choline during different gestation period alters epigenetic modifications of many imprinted genes in the genomic DNA isolated from cord blood (22, 23).

Betaine is an important methyl nutrient in animals and plants, which derived from either dietary intake or endogenous synthesis from choline (24). Betaine can improve the growth performance and muscle growth via IGF-1/mTOR signaling pathway in partridge shank broiler chickens (25). It was reported that betaine attenuates water-immersion restraint stress induced memory impairment and is regulated by the GABAergic neuronal system in the mice hippocampus (26). Also, betaine can prevents cognitive impairment and brain oxidative stress through GAT2/BGT-1 in Aβ25-35-injected mice (27). Our previous study found that maternal betaine supplementation induced intergenerational impacts on IGF1 expression and DNA methylation in offspring hepatic of F1 and F2 rats (28). Additionally, maternal betaine supplementation upregulated the expression of IGF2 through modified DNA methylation of DMRs in hippocampus of newborn piglets (29).

Epigenetic transgenerational inheritance is defined as the phenotypic change across generations and germline transmission of epigenetic information without direct genetic manipulation (30) or environmental exposure (31). A number of previous studies have shown environmental factors such as vinclozolin (30), pesticide (32) and dioxin (33) promote the epigenetic transgenerational inheritance of adult onset disease and sperm epimutations. Additionally, maternal and paternal diet also has been shown to have transgenerational behavioral and physiological effects. Paternal high-fat diet causes subfertility in both male and female offspring and grandoffspring of mice (34). Paternal folate deficiency decreases fetal brain DNA methylation levels and IGF-2 expression levels (35). Moreover, our previous study found that maternal betaine supplementation enhanced DNA methylation in IGF2/H19 ICR (36) and decreased DNA methylation of IGF-1 promoter (28) in F2 generation of rat offspring liver. Furthermore, ancestral 2,3,7,8-Tetrachlorodibenzo-p-dioxin (TCDD) exposure may promote epigenetic transgenerational alterations of imprinted gene IGF-2 in adult somatic tissue (37). However, the effect of maternal betaine supplementation on transgenerational inheritance of imprinted gene IGF-2 in offspring hippocampus is still unknown.

Therefore, the aim of this work was to investigate whether maternal betaine supplementation during pregnancy and lactation can exert transgenerational inheritance effects on estrogen-responsive IGF-2/IGFBP2 genes in hippocampus across two generations. To explore the underlying mechanisms, we determined IGF-2/IGFBP2 and ERα/β expression, as well as promoter methylation in F0, F1and F2 generation of rat hippocampus. Increased IGF-2/IGFBP2 and ERα/β expression in F0 and F2 hippocampal, accompanied by hypermethylation in IGF-2 DMRs and hypomethylation in IGF-2/ IGFBP2 and ERα/β promoters. Conversely, maternal betaine supplementation decreased IGF-2/IGFBP2 and ERα/β expression in F1 hippocampal, together with hypomethylation in IGF-2 DMRs and hypermethylation in IGF-2/ IGFBP2 and ERα/β promoters.

## 2. Results

### 2.1. Effects of maternal betaine supplementation on body weight in F1 and F2 generations

Maternal betaine supplementation significantly increased litter size (*P* < 0.05) and litter weight (*P* < 0.05) in F1 offspring, yet not influence in F2 offspring. However, maternal betaine supplementation significantly decreased birth weight (*P* < 0.01) and weaning weight (*P* < 0.01) in F1 offspring, but increased (*P* < 0.05) in F2 offspring (Table 1).

**Table 1.** Reproductive performance and body weight of female offspring

| Parameter | F1 generation |  |  | F2 generation |  |  |
| --- | --- | --- | --- | --- | --- | --- |
|  | Control | Betaine | *p* value | Control | Betaine | *p* value |
| Litter size | 13.50 ± 0.74<br>(N = 12) | 15.45 ± 0.53<br>(N = 11) | 0.047 | 15.38 ± 0.25<br>(N = 13) | 14.65 ± 0.46<br>(N = 12) | 0.155 |
| Litter weight (g) | 90.92 ± 4.00<br>(N = 12) | 101.98 ± 3.34<br>(N = 11) | 0.048 | 103.02 ± 3.42<br>(N = 13) | 106.55 ± 3.13<br>(N = 12) | 0.512 |
| Birth weight (g) | 6.74 ± 0.07<br>(n = 102) | 6.42 ± 0.08<br>(n = 128) | 0.008 | 6.50 ± 0.05<br>(n = 116) | 6.76 ± 0.07<br>(n = 101) | 0.012 |
| Weaning weight (g) | 65.58 ± 0.45<br>(n = 60) | 60.82 ± 0.68<br>(n = 60) | 0.007 | 68.51 ± 0.53<br>(n = 40) | 72.63 ± 0.75<br>(n = 40) | 0.031 |

Values are means ± SEM. “N” represents number of pregnant rats or the number of litters.
“n” represents the number of offspring rats.

### 2.2. Effects of maternal betaine supplementation on IGF-2 expression and the ICR methylation level in F0, F1 and F2 generations of rat offspring hippocampus

The structure of the IGF2/H19 locus is schematically shown in Figure 1A. Four specific primer pairs are designed to amplify 4 segments containing a motif (CCGCNNGGNGGCAG) of CTCF binding sites on the promoter of rat H19 gene (21). Maternal betaine supplementation significantly increased IGF-2 expression in F0 (*P* < 0.01), F2 (*P* < 0.05) generation (Figure 1B and 1H). These results may be related to the hypermethylation levels (*P* < 0.05) in its DMR region (Figure 1C and 1I). Simultaneously, IGF-2 protein expressions were also significantly increased in betaine group of F0 (*P* < 0.01), F2 (*P* < 0.05) generation (Figure 1D and 1J). Interestingly, in F1 generation, IGF-2 mRNA (*P* < 0.01) (Figure 1E) and protein (*P* < 0.05) (Figure 1G) expression were significantly decreased in betaine supplementation group, associated with hypomethylation levels (*P* < 0.05) in its DMR region (Figure 1F).

**Figure 1.**
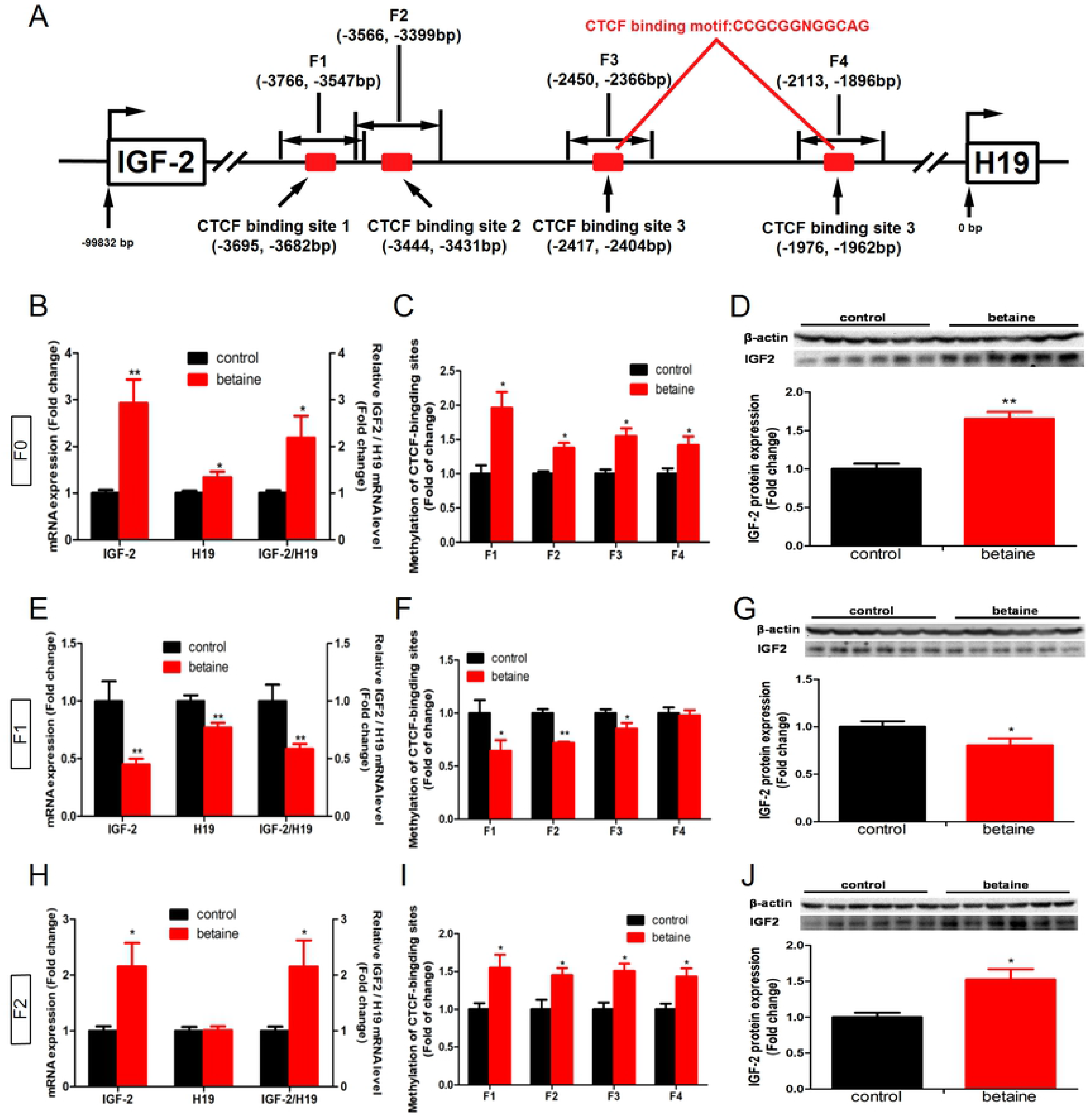
Effect of maternal betaine supplementation during pregnancy and lactation on IGF-2/H19 expression, and the methylation levels of ICR in F0, F1 and F2 generation rat offspring hippocampus. (A) The schema of designed MeDIP primers for detecting DNA methylation of four CTCF-binding sites in IGF2/H19 imprinting control region; (B); IGF2 and H19 mRNA expression in F0 generation; (C) DNA methylation levels of four CTCF-binding sites in IGF2/H19 imprinting control region in F0 generation; (D) IGF2 protein expression in F0 generation; (E); IGF2 and H19 mRNA expression in F1 generation; (F) DNA methylation levels of four CTCF-binding sites in IGF2/H19 imprinting control region in F1 generation; (G) IGF2 protein expression in F1 generation; (H); IGF2 and H19 mRNA expression in F2 generation; (I) DNA methylation levels of four CTCF-binding sites in IGF2/H19 imprinting control region in F2 generation; (J) IGF2 protein expression in F2 generation. Values are means ± SEM. \**P* < 0.05, \*\**P* < 0.01, compared with control (n = 6).

### 2.3. Effects of maternal betaine supplementation on IGF-2 mRNA variants expression and the promoter methylation level in F0, F1 and F2 generations of rat offspring hippocampus

Maternal betaine supplementation significantly increased (*P* < 0.05) IGF-2 mRNA variants expression (IGF-2 p201, p202, p203, p204) in F0, F2 generation (Figure 2B and 2F). Meanwhile, the hypomethylation levels (*P* < 0.05) were detected in IGF-2 p201 and IGF-2 p202 promoter of F0 generation (Figure 2C), and IGF-2 p201 and IGF-2 p203 promoter of F2 generation (Figure 2G). Conversely, maternal betaine supplementation significantly decreased (*P* < 0.05) IGF-2 mRNA variants expression (Figure 2D), associated with hypomethylation levels (*P* < 0.05) in IGF-2 p202, IGF-2 p203 and IGF-2 p204 promoter (Figure 2E) in F1 generation.

**Figure 2.**
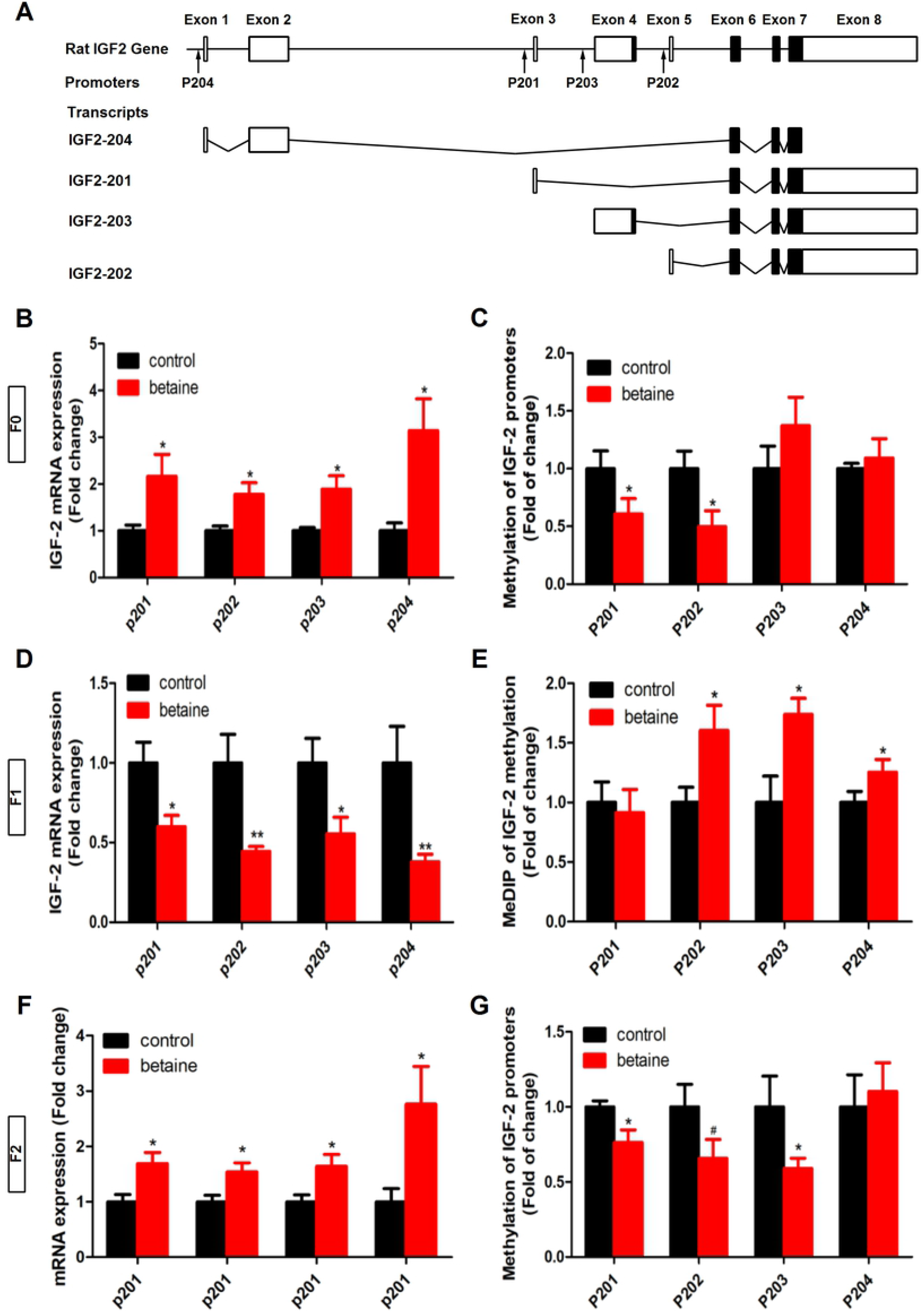
Effect of maternal betaine supplementation during pregnancy and lactation on IGF-2 mRNA variants expression, and the promoter methylation levels in F0, F1 and F2 generation rat offspring hippocampus. (A) The schema of designed MeDIP primers for detecting DNA methylation of four promoters; (B) Four IGF-2 mRNA variants expression F0 generation; (C) Four IGF-2 mRNA variants promoter methylation levels F0 generation; (D) Four IGF-2 mRNA variants expression F1 generation; (E) Four IGF-2 mRNA variants promoter methylation levels F1 generation; (F) Four IGF-2 mRNA variants expression F2 generation; (G) Four IGF-2 mRNA variants promoter methylation levels F2 generation. Values are means ± SEM. \**P* < 0.05, \*\**P* < 0.01, compared with control (n = 6).

### 2.4. Effects of maternal betaine supplementation on IGFBPs expressions and the promoter methylation levels in F0, F1 and F2 generations of rat offspring hippocampus

We detected the mRNA expression and promoter methylation levels of five IGFBPs in the hippocampus of F0, F1 and F2 rat offspring. IGFBP2 and IGFBP5 mRNA expression were significantly increased in betaine supplementation group. Maternal betaine supplementation significantly increased (*P* < 0.05) IGFBP2 mRNA expression in F0 and F2 generation (Figure 3A and 3E), accompanying by hypomethylation (*P* < 0.05) on its promoter (Figure 3B and 3F). In contrast, in F1 generation, IGFBP2 mRNA expression was significantly decreased (*P* < 0.05) in betaine supplementation group (Figure 3C), associated with hypermethylation levels (*P* < 0.01) in its promoter (Figure 3D).

**Figure 3.**
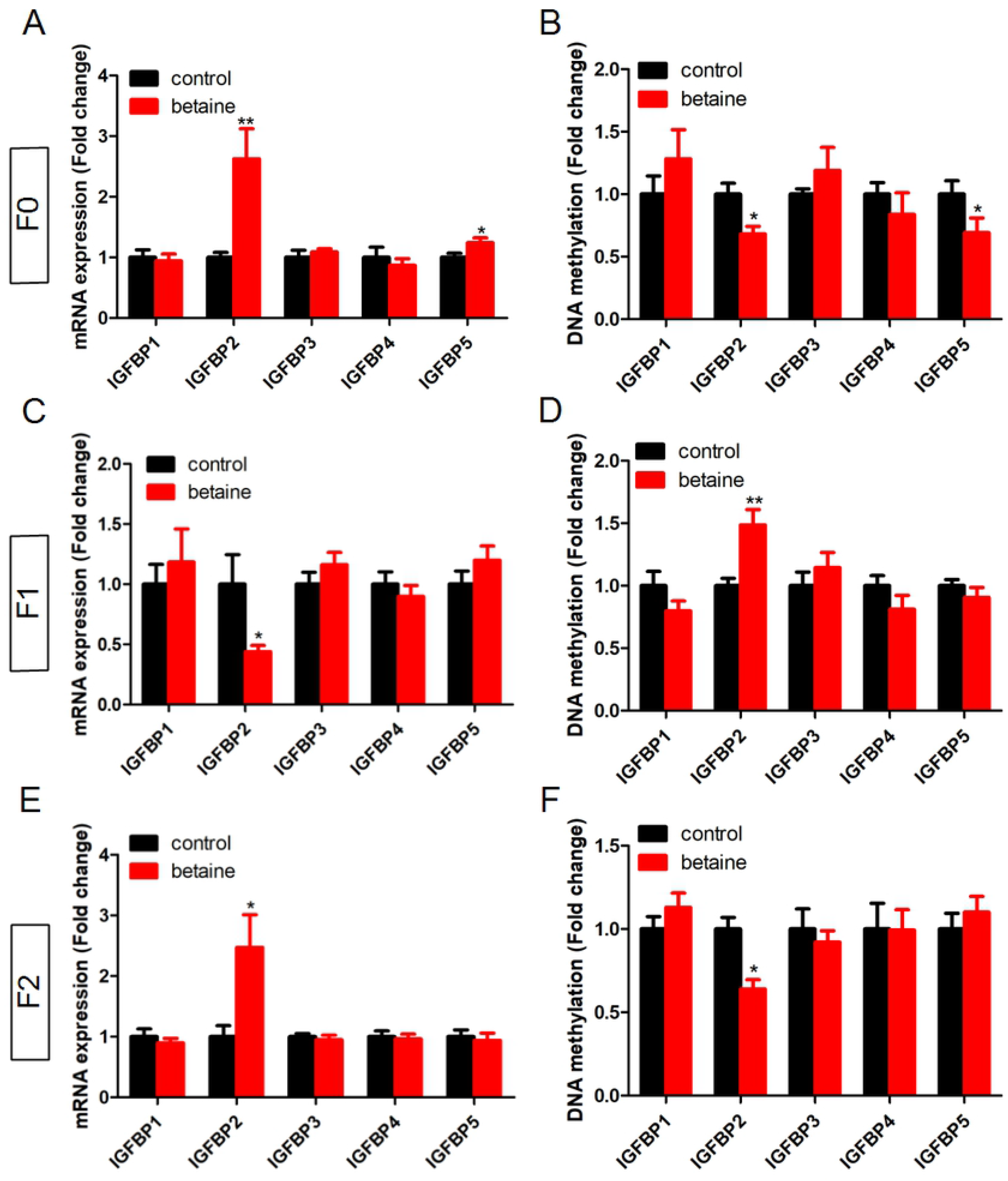
Effect of maternal betaine supplementation during pregnancy and lactation on five IGFBPs mRNA expressions, and the promoter methylation levels in F0, F1 and F2 generation rat offspring hippocampus. (A) Five IGFBPs mRNA expressions in F0 generation; (B) Five IGFBPs mRNA promoter methylation levels F0 generation; (C) Five IGFBPs mRNA expressions in F1 generation; (D) Five IGFBPs mRNA promoter methylation levels F1 generation; (E) Five IGFBPs mRNA expressions in F2 generation; (F) Five IGFBPs mRNA promoter methylation levels F2 generation. Values are means ± SEM, \**P* < 0.05, \*\**P* < 0.01, compared with control (n = 6).

### 2.5. Effects of maternal betaine supplementation on CYP19A1 and estrogen receptor mRNA expression, and the promoter methylation levels in F0, F1 and F2 generations of rat offspring hippocampus

In F0 generation, maternal betaine supplementation significantly increased (*P* < 0.05) CYP19A1, ERα and estrogen related receptor β (ERRβ) mRNA expression (Figure 4A). Which were accompanied by hypomethylation (*P* < 0.05) on its promoter, while the promoter methylation level of CYP19A1 was trend to decreased (Figure 4B). Additionally, CYP19A1 protein expression was also significantly increased (*P* < 0.01) in betaine supplementation group (Figure 4C). Similar to F0 generation, ERRβ mRNA expression was significantly increased (*P* < 0.01) and CYP19A1, ERα and ERRγ were trend to up-regulated in F2 generation (Figure 4G). At the same time, the promoter methylation level of CYP19A1, ERβ and ERRβ were significantly decreased (*P* < 0.05), and the promoter methylation level of ERα was trend to decreased in F2 generation (Figure 4H). Moreover, CYP19A1 protein expression was also significantly increased (*P* < 0.05) in betaine supplementation group (Figure 4I). Contrary to F0 and F2, in F1 generation, mRNA expression of CYP19A1, ERα, ERβ and ERRβ were significantly decreased (*P* < 0.05) in betaine supplementation group (Figure 4D). While the promoter methylation level of CYP19A1, ERα and ERRβ were significantly increased (*P* < 0.05) (Figure 4E). Furthermore, CYP19A1 protein expression was trend to down-regulated in betaine supplementation group (Figure 4F).

**Figure 4.**
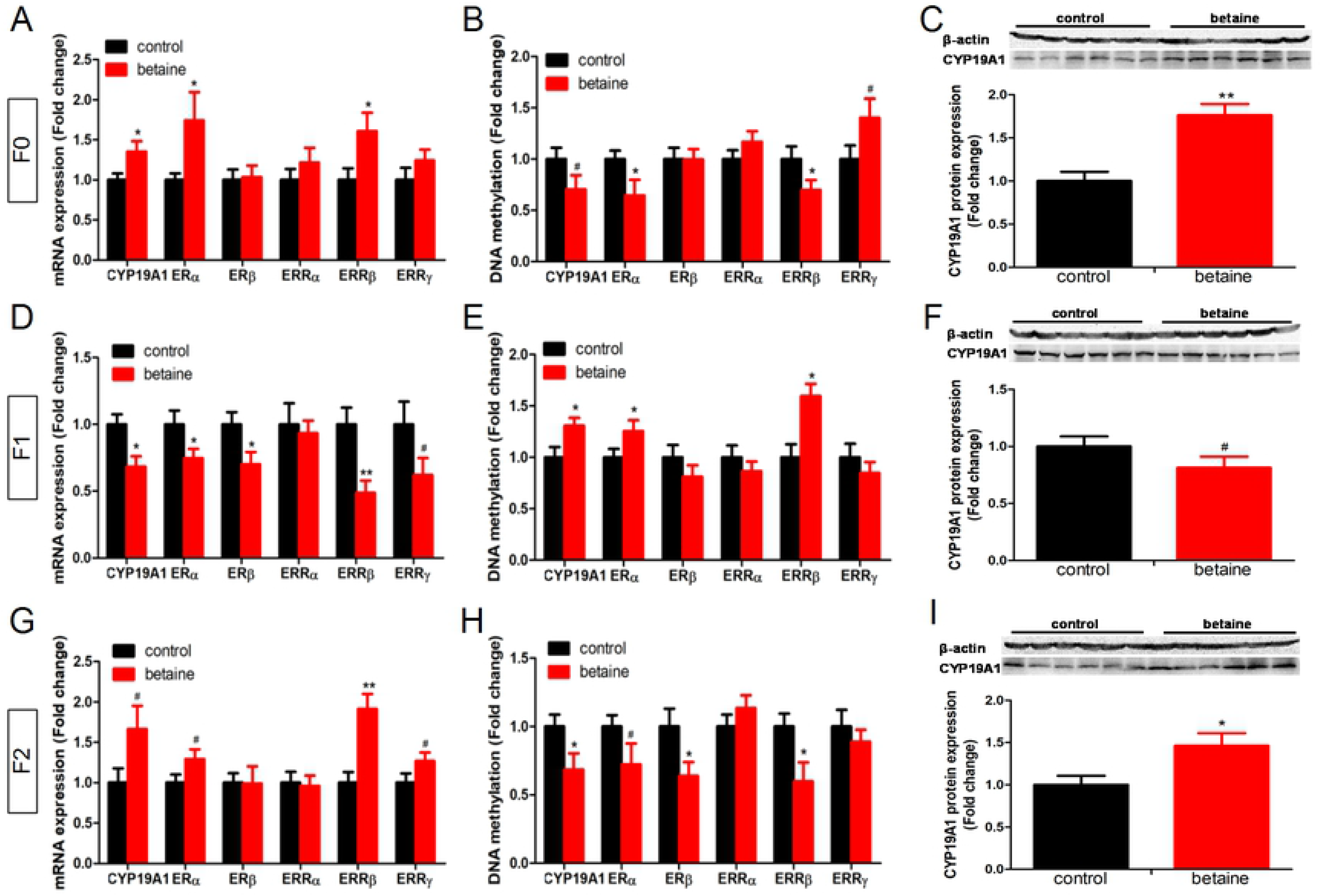
Effect of maternal betaine supplementation during pregnancy and lactation on aromatase (CYP19A1) and estrogen receptor related gene (ERα, ERβ, ERRα, ERRβ and ERRγ) mRNA expressions, and the promoter methylation levels in F0, F1 and F2 generation rat offspring hippocampus. (A) CYP19A1 and estrogen receptor related gene mRNA expressions in F0 generation; (B) CYP19A1 and estrogen receptor related gene promoter methylation levels in F0 generation; (C) CYP19A1 protein expression in F0 generation; (D) CYP19A1 and estrogen receptor related gene mRNA expressions in F1 generation; (E) CYP19A1 and estrogen receptor related gene promoter methylation levels F1 generation; (F) CYP19A1 protein expression in F1 generation; (G) CYP19A1 and estrogen receptor related gene mRNA expressions in F2 generation; (H) CYP19A1 and estrogen receptor related gene promoter methylation levels F2 generation; (I) CYP19A1 protein expression in F2 generation. Values are means ± SEM, \**P* < 0.05, \*\**P* <0.01, compared with control (n = 6).

### 2.6. Effects of maternal betaine supplementation on methionine metabolic and DNA methyltransferase related mRNA and protein expression in F0, F1 and F2 generation hippocampus

In F0 generation, maternal betaine supplementation significantly increased (*P* < 0.05) BHMT, AHCY, DNMT1 and DNMT3A mRNA expression (Figure 5A), and BHMT protein expression in betaine supplementation group (Figure 5B). Meanwhile, maternal betaine supplementation significantly increased (*P* < 0.05) BHMT mRNA and protein expression in F2 generation (Figure 5E and 5F). Conversely, maternal betaine supplementation significantly decreased (*P* < 0.05) BHMT, AHCY and DNMT3A mRNA expression (Figure 5A) and BHMT protein expression in betaine supplementation group (Figure 5B).

**Figure 5.**
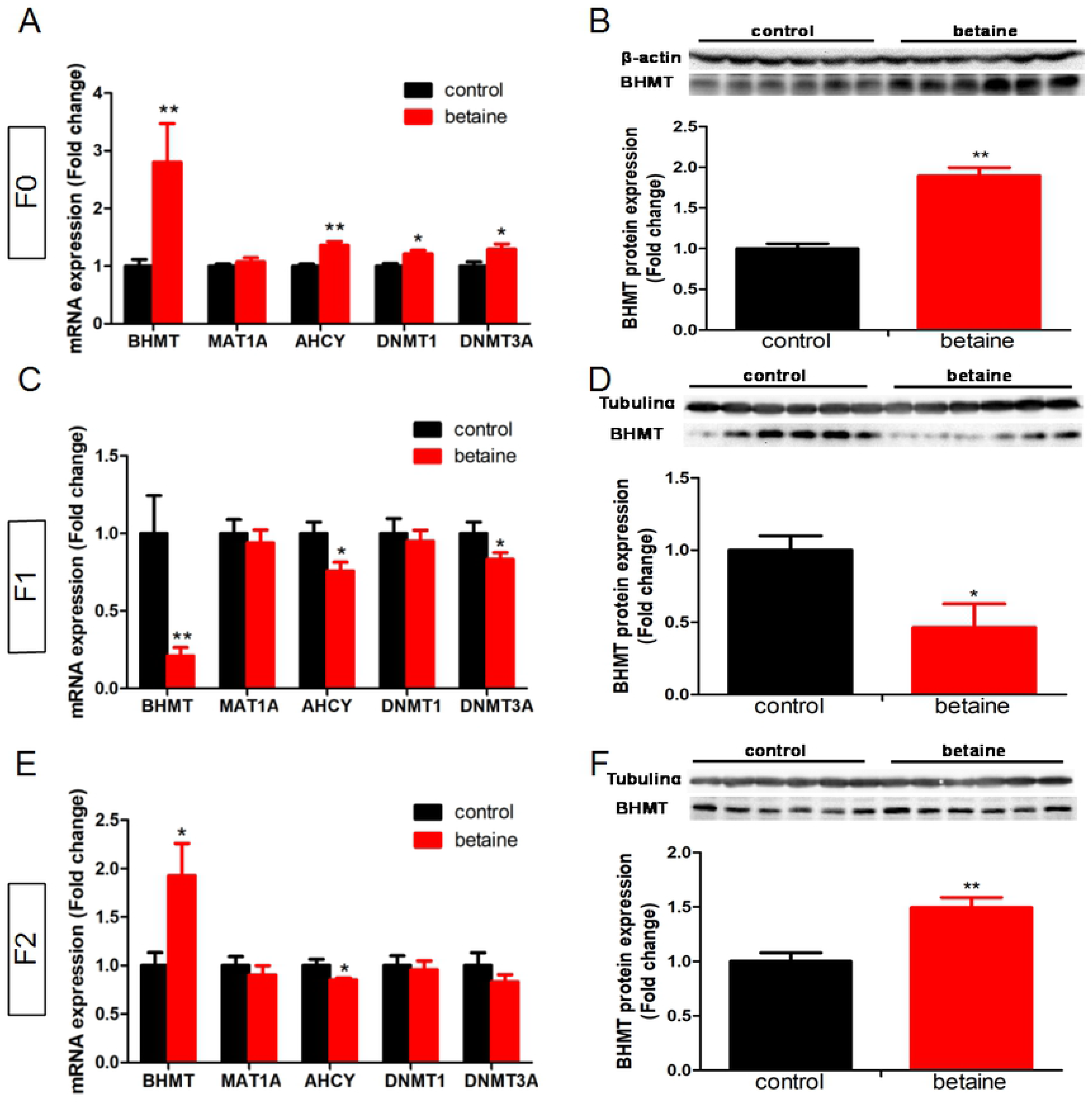
Effect of maternal betaine supplementation during pregnancy and lactation on methionine metabolism and DNA methyltransferase related gene mRNA and expressions in F0, F1 and F2 generation rat offspring hippocampus. (A) BHMT, MAT1A, AHCY, DNMT1 and DNMT3A mRNA expressions in F0 generation; BHMT protein expression in F0 generation; (C) BHMT, MAT1A, AHCY, DNMT1 and DNMT3A mRNA expressions in F1 generation; (D) BHMT protein expression in F1 generation; (E) BHMT, MAT1A, AHCY, DNMT1 and DNMT3A mRNA expressions in F2 generation; (F) BHMT protein expression in F2 generation. Values are means ± SEM, n =6. alues are means ± SEM, \**P* < 0.05, \*\**P* <0.01, compared with control (n = 6).

## 3. Discussion

The current study demonstrated the maternal methyl nutrition betaine exerts an epigenetic transgenerational effect on hippocampal expression of estrogen-responsive genes in rat offspring, which is associated with corresponding alterations in DNA methylation on ICR of IGF-2 gene and the promoter of affected genes. Previous studies shown that methyl donor supplementation increased the fetal weight (40) and birth weight (41) in pigs. Similar, maternal methyl donor restriction decreased the birth weight in rats (42). However, in the present study, maternal betaine supplementation induced retarded growth and decreased body weight in F1 generation. This result is in line with a previous finding that an excess methyl donor supplementation in the perinatal period reduced the body weight of female rat offspring (43). Interestingly, we found that maternal betaine supplementation in F0 generation increased the birth and weaning weight of F2 offspring. Similar transgenerational inheritance effects were reported that during Dutch famine period, F1 offspring derived from starved mother during pregnancy had lower birth weight, yet F2 offspring had higher adult body weight (44). Additionally, F0 generation gestating female rats exposed to dichlorodiphenyltrichloroethane (DDT) were not found to develop obesity in F1 offspring, whereas in F3 generation (great grand-offspring) had over 50% of males and females develop obesity (45). Although complex interplay among environmental, metabolic and epigenetic factors has been proposed to participate in such nongenetic transgenerational inheritance (46, 47), the mechanisms underlying the multigenerational transmission of phenotypes are not fully understood. Furthermore, we found that F1 litter size and litter weight were increased in betaine-supplemented group. This result is also against the previous reports that maternal methyl donor supplementation or restriction did not affect the litter size in pigs (41), even reduced litter size in rats (42). Above these inconsistent studies may be explained by differences in the animal species and diet compositions, the types and levels of methyl donors, as well as the time and period of supplementation.

Fetal growth is largely controlled by the complex insulin/IGF system. In particular, several growth-related imprinted genes are related to the expression and function of the potent fetal growth factor, like IGF-2 (48). Loss of imprinting of IGF-2 results in biallelic expression, excess production of IGF-2 transcripts and increased levels of IGF-2 protein. Also, disruptions at imprinted gene loci, such as IGF-2 and H19 have been demonstrated to significantly disrupt allometric growth and development of specific relevant organs including heart, liver and kidney (48, 49). This could directly result in adult onset diseases or manifest in abnormal phenotype due to pleiotropic effects of the gene imprinted in the fetus (50). Conversely, previous study has been demonstrated that imprinted gene loci IGF-2 can promote growth and development of specific relevant organs such as liver (51) and hippocampus (13). In this study, we found that hippocampus IGF-2 and its four mRNA variants (p201, p202, p203 and p204) were up-regulated in maternal betaine supplementation group, and these effects were still manifested in the F2 generation. Surprisingly, this effect skips F1 generation, even IGF-2 and its four mRNA variants were decreased in F1 generation. This result is against the previous reports that ancestral p,p’-dichlorodiphenoxydichloroethylene (p,p’-DDE) exposure reduced IGF-2 expression in three successive generations, from F1 to F3 generation (52). This inconsistent study may be explained by differences in the animal species and exposure compositions. Meanwhile, because of IGF-2 mRNA and protein increased in F0 and F2 generation, whereas decreased in F1 generation. The body weight of pups decreased in F1 generation, but increased in F2 generation may be partly explained by the related change of IGF-2.

IGF-2 and H19 are reciprocally expressed imprinted genes which are located in the IGF-2/H19 ICR and share a downstream enhancer. ICR is located 2-4 kb upstream of the H19 transcription start site between IGF-2 and H19, as a methylation-regulated enhancer blocker to mediate the monoallelic expression of IGF-2 and H19 through four CTCF binding sites within the ICR (53). CTCF can remotely control IGF-2 through long-range chromatin interaction (53) and maintain the hypomethylation of the ICR in somatic tissues (54). Previous study reported that decreased mRNA and protein levels of IGF-2, in combination with the hypomethylation of ICR in the resorbing embryo and spermatozoa of tamoxifen-treated rats (55). In this study, maternal betaine supplementation could significantly induce hypermethylation in four CTCF binding site in the ICR of the F0 and F2 generation, whereas induce hypomethylation in three CTCF binding site in the ICR of the F1 generation. These change of methylation levels in IGF-2 ICR can be used to explain IGF-2 mRNA expression in F0, F1 and F2 generation. These results is similar to the previous reports that ancestral TCDD exposure were found to decreased imprinted gene IGF-2 expression in F1 and F3 generation, accompanied with hypomethylation of DMR2 in the liver of offspring in F1 and F3 generation (37). However, this result is also against the previous reports that paternal benzo[a]pyrene (BaP) exposure alters the sperm DNA methylation levels of imprinting genes IGF-2 in F0 generation mice and their unexposed F1-F2 male offspring (56). And, ancestral p,p’-DDE exposure reduced IGF-2 expression, in combination with the hypomethylation of DMR2 in three successive generations (F1 to F3 generation) (52). This inconsistent study may be explained by differences in the animal species, ancestral gender and tissue type, as well as exposure compositions. Consistent with these transgenerational IGF-2 epigenetic effects in above studies, the present findings indicated that maternal betaine exposure induced a transgenerational hypermethylation of IGF-2 ICR from F0 to F2 rat offspring hippocampus.

It is also noted that the intergenerational effects of grand dam betaine exposure also shown in the promoter methylation of IGF-2 mRNA variants. The decreased of DNA methylation levels on IGF-2 mRNA variants promoter in F0 and F2 generation, but increased in F1 generation. These results may be used to explain the related change of IGF-2 mRNA variants expression. Furthermore, the intergenerational effects of grand dam betaine exposure appear to be gene specific. Hippocampus IGFBP2 mRNA expression was modified, being consistent with IGF-2, in response to betaine supplementation. It was significantly upregulated in F0 and F2, but downregulated in F1 pups. Meanwhile, related changes in mRNA expression were associated with accordant alterations in promoter methylation. Previous study has evidence that IGFBP-2 is abundant in brain and is highly expressed by astrocytes in the cortex, and plays an important role in the nervous development (57). IGFBP2 also has the ability to bind insulin, IGF1 and IGF2 with an increased affinity for the latter (58). IGF-2/IGFBP2 complex was shown that have a marked affinity for extra cellular matrix (59), and also was speculate that may stimulate IGF signaling in response to estrogen in rat hippocampus (12). In this study, we surprisingly found that IGF-2 and IGFBP2 expression have the same mode in three generation, not only mRNA expression, but also the promoter methylation levels. Thus, we speculate that maternal betaine exposure induce the change of body weight in offspring may be associated with the related change of IGF-2/IGFBP2 complex expression.

Interestingly, aromatase (CYP19A1), ERα and estrogen related receptor β (ERRβ) were modified in three generation, being opposite to their promoter methylation levels, and even also have the intergenerational effects induced by maternal betaine exposure in rat hippocampus. Thus, these changes in mRNA levels may be related to their promoter hypo/hypermethylation levels. Similar study has been reported that maternal folic acid supplementation during gestation reduces ERα expression in rat offspring (60). In contrast, early methyl donor deficiency reduces aromatase, ERα and ERβ expression in the cerebellum of female rat pups (61). Methyl donor nutrition vitamin B12 and folate during pregnancy and postnatal life impairs proper brain development via impaired influence of ERα (62). Neuronal aromatase activity may play an important role in the putative neuroprotective action of estrogens. Estrogens has been reported that not only increase spine and synapse density in CA1 of the hippocampus and to impact hippocampal synaptic plasticity, but also stimulate proliferation and differentiation of progenitors into mature granule neurons in the DG (63, 64). These effects of estrogen in the hippocampus may be mediated by ERα and ERβ, which widely distributed and expressed in the hippocampus, cerebral cortex, amygdale and etc., these regions helps to regulate the autonomic nervous system, memory and cognition, and emotional reactions (65, 66). In our investigations, CYP19A1, ERα and ERRβ expression were increased in F0 generation and trend to increase in F2 generation, but decreased in F2 generation in maternal betaine exposure rat offspring hippocampus. These results indicated that maternal betaine supplementation may affect hippocampus neurogenesis in F0 to F3 rat offspring. Furthermore, IGF-2 and IGFBP2 were found to respond to estrogen in hippocampus of young female rats (12). Thus, Thus, combined with changes in IGF-2 and IGFBP2 in present study, we speculate that maternal betaine exposure induce the change of body weight in offspring may be associated with the related change of estrogen response gene IGF-2/IGFBP2 complex expression.

Overall, our study provides evidence that ancestral exposure to betaine may promote epigenetic transgenerational inheritance of IGF-2/IGFBP2, aromatase and ER expression from F0 to F2 generation, accompanied with promoter DNA methylation, and even leads to distinct growth phenotype in F1 and F2 offspring. Nevertheless, these results in present study may provide and valuable clues for the mechanism of maternal methyl donor nutrition influence the brain function and growth performance in F1 and F2 offspring. Undoubtedly, further research is necessary to elucidate the interplay between intergenerational epigenetics and transgenerational inheritance.

## 4. Materials and Methods

### 4.1 Ethics statement

The experimental protocol was approved by the Animal Ethics Committee of Nanjing Agricultural University, with a project number 2016YFD0500502. The sampling procedures complied with the “Guidelines on Ethical Treatment of Experimental Animals” (2006) No.398 set by the Ministry of Science and Technology, China.

### 4.2 Animals and experimental design

Three-month-old Sprague-Dawley female rats weighing 400 g in average were purchased from the Laboratory Animal Center of Jiangsu University. All rats were housed in a controlled temperature (22 ± 0.5°C) and humidity (50 ± 5%) with artificial lighting (12-hour light/12-hour dark cycle). The animals had free access to both food and water. All together 40 female rats (F0) were mated during a period of 3 weeks. Two female rats were housed with a male and fed with basal diet. Mating was confirmed by the presence of a vaginal smear plug. Mated F0 female rats were transferred to individual cages and randomly divided into two groups. The control group fed a basal diet and betaine group fed a diet supplemented with betaine (98% purity, B2629, Sigma Chemical Co., St. Louis, MO, USA) at the level of 10 g/kg according to previous publications (38, 39). Both basal and betaine-supplemented diets were custom-made by Jiangsu Xie Tong Feed Company (Nanjing, Jiangsu, China). After birth (D0), litter size was adjusted to 10 pups (5 males and 5 females). Randomly selected 20 weaned (D21) pups of F1 females fed with basal diet thereafter from control and betaine group and others killed with pentobarbital sodium. At 3 months of age, F1 females were mated with males that received basal diet throughout life. According to the dietary treatment of their grandmothers, the derived F2 offspring rats were categorized to control and betaine groups, respectively. During the pregnancy and lactation, F1 females were fed basal diet. Litter size was adjusted to 10 at birth. F2 pups were weaned at day 21 (D21) and 20 female F2 rats weaned on the same day were killed with pentobarbital sodium.

### 4.3 RNA isolation and real-time PCR

Total RNA was isolated from 30 mg hippocampus samples using 600 μL TRIzol reagents (Invitrogen, Carlsbad, California, USA). One microgram of RNA samples was reverse-transcribed according to the manufacturer’s protocol (Vazyme Biotech, Nanjing, Jiangsu, China). One μL of diluted cDNA (1:25) was used for real-time PCR in a QuantStuioTM 6 Flex Real-Time PCR System (Applied Biosystems, Foster City, California, USA). Peptidylprolyl isomerase A (PPIA) was used as an internal control to normalize the technical variations. Data were analyzed using the method of 2-ΔΔCT and presented relative to the control group. All primers (Table 2) were synthesized by Suzhou GENEWIZ Biological Technology Co., Ltd (Suzhou, Jiangsu, China).

**Table 2.**
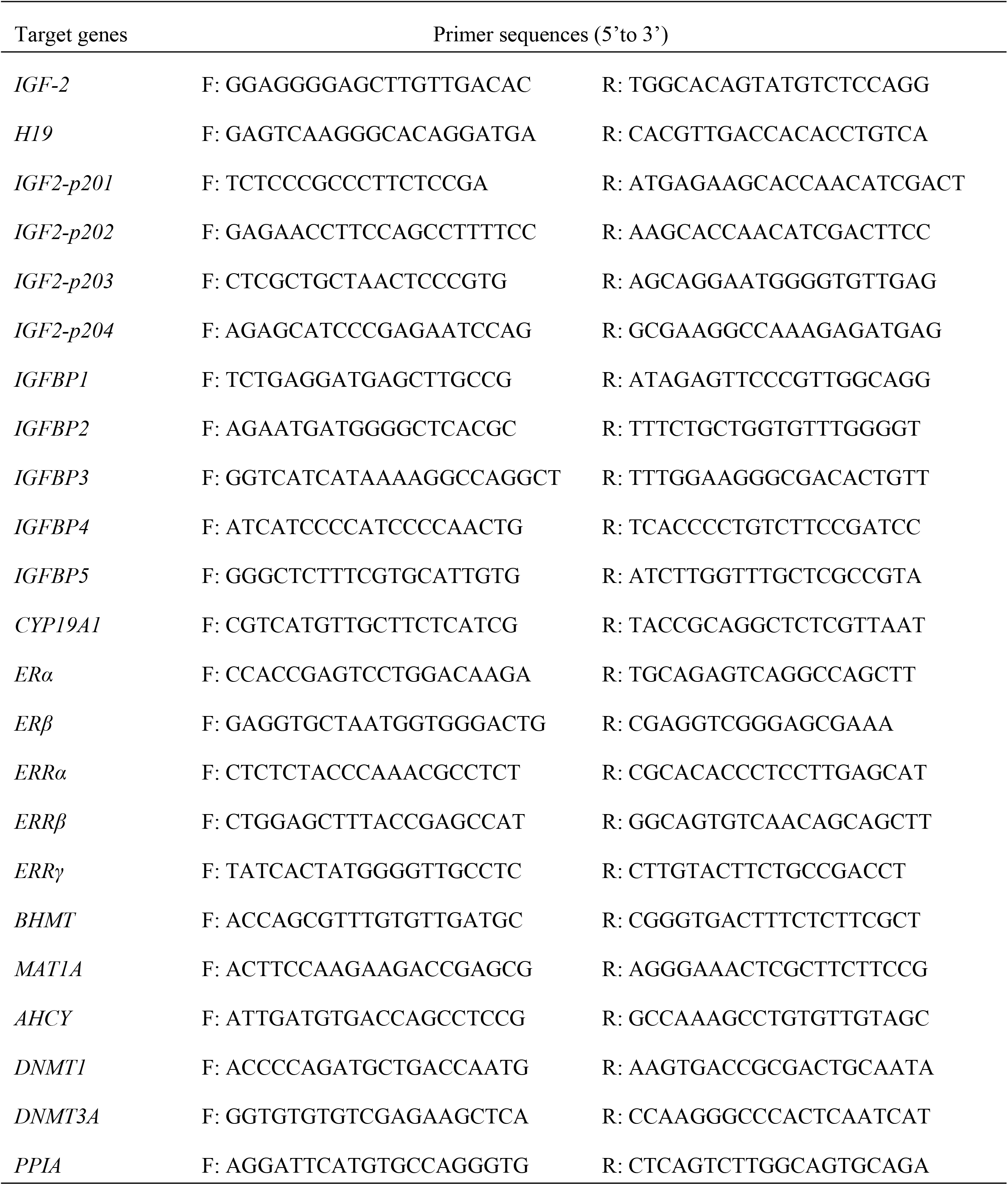
The primers sequences for RT-PCR

### 4.4 Total Protein Extraction and Western Blotting

Total protein was extracted from 50 mg frozen hippocampus samples as previously described. Protein concentrations were measured using BCA Protein Assay kit (NO.23227, Thermo Scientific, Rockford, Illinois, USA) according to the manufacturer’s instructions. Protein (60 μg/lane) was loaded for electrophoresis on a 6-14% SDS-PAGE gel and transferred onto a nitrocellulose membrane. After transfer, the membranes were blocked with 4% milk and then incubated with primary and secondary antibodies. Western blot analysis for IGF2 (BS6632, Bioworld, diluted 1:1000), CYP19A1 (BS6580, Bioworld, diluted 1:1000), betaine homocysteine methyltransferase (BHMT) (15965-1-AP, Proteintech, diluted 1:500), was carried out according to the recommended protocols provided by the manufacturers, and β-actin (AC026, ABclonal, diluted 1:50,000), Tubulinα (BS1699, Bioworld, diluted 1:10,000) was used as loading control. Images were captured by VersaDoc 4000MP system (Bio-Rad, USA) and the band density was analyzed with Quantity One software (Bio-Rad, USA).

### 4.5 Methylated DNA immunoprecipitation (MeDIP)

High-quality genomic DNA was extracted from hippocampal and sonicated to fragments of approximately 500 bp in size. Two micrograms of sonicated DNA were heat-denatured to produce single-stranded DNA, and a portion of the sonicated DNA was stored as control (input) DNA. A mouse monoclonal antibody against 5-methyl cytidine (AB10805, Abcam, UK) was used to immunoprecipitate methylated DNA fragments. The immune complexes were captured with protein G agarose beads pretreated with denatured salmon sperm DNA and BSA (No. P2009, Beyotime Institute of Biotechnology). The beads were washed to eliminate nonspecific binding and resuspended in 250 μL digestion buffer containing proteinase K. Finally, the MeDIP DNA was purified. Finally, the MeDIP DNA was purified, diluted 1/25 and used to amplify the sequences of CTCF binding sites of IGF2/H19 ICR and IGF2, IGFBPs gene promoters by real-time PCR with specific primers (Table 3).

**Table 3.**
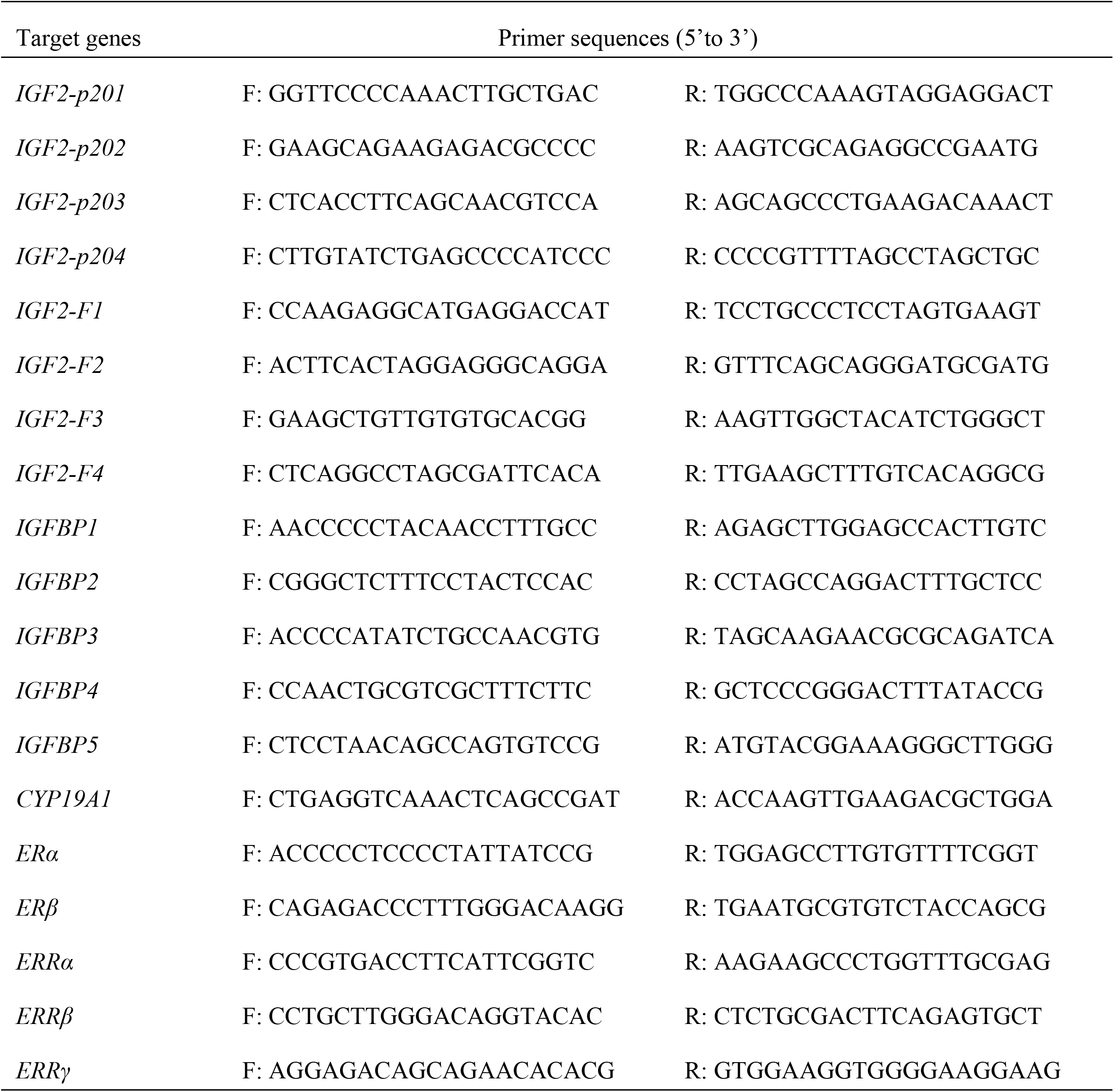
The primers sequences for MeDIP assay

### 4.6. Statistical analysis

All data are presented as means ± SEM and the differences among groups were analyzed using independent-samples T-Test with SPSS 20.0 for windows. The differences were considered statistically significant when P < 0.05.

## Acknowledgment

This work was supported by the National Key Research and Development Program of China (2016YFD0500502), the National Basic Research Program of China (2012CB124703), the Fundamental Research Funds for the Central Universities (KYZ201212), the Priority Academic Program Development of Jiangsu Higher Education Institutions (PAPD), the Postgraduate Research & Practice Innovation Program of Jiangsu Province (KYCX18_0716), and Jiangsu Collaborative Innovation Center of Meat Production and Processing, Quality, and Safety Control. The funders had no role in study design, data collection and analysis, decision to publish, or preparation of the manuscript.

## References

1. Bartsch T, Wulff P. The hippocampus in aging and disease: From plasticity to vulnerability. Neuroscience. 2015 Nov 19;309:1–16. PubMed PMID: 26241337.

2. Spencer JL, Waters EM, Romeo RD, Wood GE, Milner TA, McEwen BS. Uncovering the mechanisms of estrogen effects on hippocampal function. Frontiers in neuroendocrinology. 2008 May;29(2):219–37. PubMed PMID: 18078984. PubMed Central PMCID: 2440702.

3. Foy MR. Ovarian hormones, aging and stress on hippocampal synaptic plasticity. Neurobiology of learning and memory. 2011 Feb;95(2):134–44. PubMed PMID: 21081173. PubMed Central PMCID: 3045646.

4. Kretz O, Fester L, Wehrenberg U, Zhou LP, Brauckmann S, Zhao ST, et al. Hippocampal synapses depend on hippocampal estrogen synthesis. J Neurosci. 2004 Jun 30;24(26):5913–21. PubMed PMID: WOS:000222351900008. English.

5. Prange-Kiel J, Rune GM. Direct and indirect effects of estrogen on rat hippocampus. Neuroscience. 2006;138(3):765–72. PubMed PMID: 16324798.

6. Vierk R, Glassmeier G, Zhou LP, Brandt N, Fester L, Dudzinski D, et al. Aromatase Inhibition Abolishes LTP Generation in Female But Not in Male Mice. J Neurosci. 2012 Jun 13;32(24):8116–26. PubMed PMID: WOS:000305295600004. English.

7. Brocca ME, Garcia-Segura LM. Non-reproductive Functions of Aromatase in the Central Nervous System Under Physiological and Pathological Conditions. Cellular and molecular neurobiology. 2019 May;39(4):473–81. PubMed PMID: 30084008.

8. Fester L, Zhou LP, Butow A, Huber C, von Lossow R, Prange-Kiel J, et al. Cholesterol-Promoted Synaptogenesis Requires the Conversion of Cholesterol to Estradiol in the Hippocampus. Hippocampus. 2009;19(8):692–705. PubMed PMID: WOS:000268988400002. English.

9. Brake WG, Alves SE, Dunlop JC, Lee SJ, Bulloch K, Allen PB, et al. Novel target sites for estrogen action in the dorsal hippocampus: An examination of synaptic proteins. Endocrinology. 2001 Mar;142(3):1284–9. PubMed PMID: WOS:000167166400040. English.

10. Prange-Kiel J, Fester L, Zhou L, Lauke H, Carretero J, Rune GM. Inhibition of hippocampal estrogen synthesis causes region-specific downregulation of synaptic protein expression in hippocampal neurons. Hippocampus. 2006;16(5):464–71. PubMed PMID: WOS:000237428900005. English.

11. Tang SS, Ren Y, Ren XQ, Cao JR, Hong H, Ji H, et al. ERalpha and/or ERbeta activation ameliorates cognitive impairment, neurogenesis and apoptosis in type 2 diabetes mellitus mice. Experimental neurology. 2019 Jan;311:33–43. PubMed PMID: 30201537.

12. Takeo C, Ikeda K, Horie-Inoue K, Inoue S. Identification of Igf2, Igfbp2 and Enpp2 as Estrogen-Responsive Genes in Rat Hippocampus. Endocr J. 2009 Feb;56(1):113–20. PubMed PMID: WOS:000264147800014. English.

13. Li CQ, Luo YW, Bi FF, Cui TT, Song L, Cao WY, et al. Development of anxiety-like behavior via hippocampal IGF-2 signaling in the offspring of parental morphine exposure: effect of enriched environment. Neuropsychopharmacology: official publication of the American College of Neuropsychopharmacology. 2014 Nov;39(12):2777–87. PubMed PMID: 24889368. PubMed Central PMCID: 4200488.

14. Couce ME, Weatherington AJ, McGinty JF. Expression of insulin-like growth factor-II (IGF-II) and IGF-II/mannose-6-phosphate receptor in the rat hippocampus: an in situ hybridization and immunocytochemical study. Endocrinology. 1992 Oct;131(4):1636–42. PubMed PMID: 1396308.

15. Chesik D, De Keyser J, Wilczak N. Insulin-like growth factor binding protein-2 as a regulator of IGF actions in CNS: Implications in multiple sclerosis. Cytokine Growth F R. 2007 Jul-Aug;18(3-4):267–78. PubMed PMID: WOS:000248074700007. English.

16. Ocrant I, Fay CT, Parmelee JT. Characterization of insulin-like growth factor binding proteins produced in the rat central nervous system. Endocrinology. 1990 Sep;127(3):1260–7. PubMed PMID: 1696881.

17. Lund J, Sondergaard MT, Conover CA, Overgaard MT. Heparin-binding mechanism of the IGF2/IGF-binding protein 2 complex. Journal of molecular endocrinology. 2014 Jun;52(3):345–55. PubMed PMID: 24604839.

18. DeChiara TM, Robertson EJ, Efstratiadis A. Parental imprinting of the mouse insulin-like growth factor II gene. Cell. 1991 Feb 22;64(4):849–59. PubMed PMID: 1997210.

19. Bartolomei MS, Zemel S, Tilghman SM. Parental imprinting of the mouse H19 gene. Nature. 1991 May 9;351(6322):153–5. PubMed PMID: 1709450.

20. Thorvaldsen JL, Duran KL, Bartolomei MS. Deletion of the H19 differentially methylated domain results in loss of imprinted expression of H19 and Igf2. Genes & development. 1998 Dec 1;12(23):3693–702. PubMed PMID: 9851976. PubMed Central PMCID: 317260.

21. Bell AC, Felsenfeld G. Methylation of a CTCF-dependent boundary controls imprinted expression of the Igf2 gene. Nature. 2000 May 25;405(6785):482–5. PubMed PMID: 10839546.

22. Pauwels S, Ghosh M, Duca RC, Bekaert B, Freson K, Huybrechts I, et al. Dietary and supplemental maternal methyl-group donor intake and cord blood DNA methylation. Epigenetics. 2017 Jan 2;12(1):1–10. PubMed PMID: 27830979. PubMed Central PMCID: 5270634.

23. Pauwels S, Ghosh M, Duca RC, Bekaert B, Freson K, Huybrechts I, et al. Maternal intake of methyl-group donors affects DNA methylation of metabolic genes in infants. Clinical epigenetics. 2017;9:16. PubMed PMID: 28191262. PubMed Central PMCID: 5297118.

24. Day CR, Kempson SA. Betaine chemistry, roles, and potential use in liver disease. Bba-Gen Subjects. 2016 Jun;1860(6):1098–106. PubMed PMID: WOS:000375165300006. English.

25. Chen R, Zhuang S, Chen YP, Cheng YF, Wen C, Zhou YM. Betaine improves the growth performance and muscle growth of partridge shank broiler chickens via altering myogenic gene expression and insulin-like growth factor-1 signaling pathway. Poultry science. 2018 Dec 1;97(12):4297–305. PubMed PMID: 30085311.

26. Kunisawa K, Kido K, Nakashima N, Matsukura T, Nabeshima T, Hiramatsu M. Betaine attenuates memory impairment after water-immersion restraint stress and is regulated by the GABAergic neuronal system in the hippocampus. Eur J Pharmacol. 2017 Feb 5;796:122–30. PubMed PMID: WOS:000393529300015. English.

27. Ibi D, Tsuchihashi A, Nomura T, Hiramatsu M. Involvement of GAT2/BGT-1 in the preventive effects of betaine on cognitive impairment and brain oxidative stress in amyloid beta peptide-injected mice. Eur J Pharmacol. 2019 Jan 5;842:57–63. PubMed PMID: WOS:000453397300007. English.

28. Zhao N, Yang S, Hu Y, Dong H, Zhao R. Maternal betaine supplementation in rats induces intergenerational changes in hepatic IGF-1 expression and DNA methylation. Molecular nutrition & food research. 2017 Aug;61(8). PubMed PMID: 28239993.

29. Li X, Sun QW, Li X, Cai DM, Sui SY, Jia YM, et al. Dietary betaine supplementation to gestational sows enhances hippocampal IGF2 expression in newborn piglets with modified DNA methylation of the differentially methylated regions. Eur J Nutr. 2015 Oct;54(7):1201–10. PubMed PMID: WOS:000361634700016. English.

30. Anway MD, Cupp AS, Uzumcu M, Skinner MK. Epigenetic transgenerational actions of endocrine disruptors and male fertility. Science. 2005 Jun 3;308(5727):1466–9. PubMed PMID: 15933200.

31. Skinner MK, Manikkam M, Guerrero-Bosagna C. Epigenetic transgenerational actions of environmental factors in disease etiology. Trends in endocrinology and metabolism: TEM. 2010 Apr;21(4):214–22. PubMed PMID: 20074974. PubMed Central PMCID: 2848884.

32. Manikkam M, Tracey R, Guerrero-Bosagna C, Skinner MK. Pesticide and insect repellent mixture (permethrin and DEET) induces epigenetic transgenerational inheritance of disease and sperm epimutations. Reproductive toxicology. 2012 Dec;34(4):708–19. PubMed PMID: 22975477. PubMed Central PMCID: 3513590.

33. Manikkam M, Tracey R, Guerrero-Bosagna C, Skinner MK. Dioxin (TCDD) induces epigenetic transgenerational inheritance of adult onset disease and sperm epimutations. PloS one. 2012;7(9):e46249. PubMed PMID: 23049995. PubMed Central PMCID: 3458876.

34. Fullston T, Palmer NO, Owens JA, Mitchell M, Bakos HW, Lane M. Diet-induced paternal obesity in the absence of diabetes diminishes the reproductive health of two subsequent generations of mice. Human reproduction. 2012 May;27(5):1391–400. PubMed PMID: 22357767.

35. Kim HW, Kim KN, Choi YJ, Chang N. Effects of paternal folate deficiency on the expression of insulin-like growth factor-2 and global DNA methylation in the fetal brain. Molecular nutrition & food research. 2013 Apr;57(4):671–6. PubMed PMID: 23229416.

36. Yang S, Zhao N, Yang Y, Hu Y, Dong H, Zhao R. Mitotically Stable Modification of DNA Methylation in IGF2/H19 Imprinting Control Region Is Associated with Activated Hepatic IGF2 Expression in Offspring Rats from Betaine-Supplemented Dams. Journal of agricultural and food chemistry. 2018 Mar 21;66(11):2704–13. PubMed PMID: 29376352.

37. Ma J, Chen X, Liu Y, Xie Q, Sun Y, Chen J, et al. Ancestral TCDD exposure promotes epigenetic transgenerational inheritance of imprinted gene Igf2: Methylation status and DNMTs. Toxicology and applied pharmacology. 2015 Dec 1;289(2):193–202. PubMed PMID: 26455773.

38. Kheradmand A, Alirezaei M, Dezfoulian O. Cadmium-Induced Oxidative Stress in the Rat Testes: Protective Effects of Betaine. Int J Pept Res Ther. 2013 Dec;19(4):337–44. PubMed PMID: WOS:000325773800008. English.

39. Yagisawa M, Doi Y, Uenohara T, Toda M, Shigematsu N, Nakata R. Betaine supplementation suppresses plasma homocysteine level elevation induced by folate deficiency in rats. Nutr Res. 2006 Jun;26(6):266–70. PubMed PMID: WOS:000240029500004. English.

40. Oster M, Nuchchanart W, Trakooljul N, Murani E, Zeyner A, Wirthgen E, et al. Methylating micronutrient supplementation during pregnancy influences foetal hepatic gene expression and IGF signalling and increases foetal weight. Eur J Nutr. 2016 Jun;55(4):1717–27. PubMed PMID: 26169873.

41. Jia Y, Song H, Gao G, Cai D, Yang X, Zhao R. Maternal Betaine Supplementation during Gestation Enhances Expression of mtDNA-Encoded Genes through D-Loop DNA Hypomethylation in the Skeletal Muscle of Newborn Piglets. Journal of agricultural and food chemistry. 2015 Nov 25;63(46):10152–60. PubMed PMID: 26527363.

42. Konycheva G, Dziadek MA, Ferguson LR, Krageloh CU, Coolen MW, Davison M, et al. Dietary methyl donor deficiency during pregnancy in rats shapes learning and anxiety in offspring. Nutr Res. 2011 Oct;31(10):790–804. PubMed PMID: 22074804.

43. Giudicelli F, Brabant AL, Grit I, Parnet P, Amarger V. Excess of methyl donor in the perinatal period reduces postnatal leptin secretion in rat and interacts with the effect of protein content in diet. PloS one. 2013;8(7):e68268. PubMed PMID: 23840890. PubMed Central PMCID: 3698130.

44. Veenendaal MV, Painter RC, de Rooij SR, Bossuyt PM, van der Post JA, Gluckman PD, et al. Transgenerational effects of prenatal exposure to the 1944-45 Dutch famine. BJOG: an international journal of obstetrics and gynaecology. 2013 Apr;120(5):548–53. PubMed PMID: 23346894.

45. Skinner MK, Manikkam M, Tracey R, Guerrero-Bosagna C, Haque M, Nilsson EE. Ancestral dichlorodiphenyltrichloroethane (DDT) exposure promotes epigenetic transgenerational inheritance of obesity. BMC medicine. 2013 Oct 23;11:228. PubMed PMID: 24228800. PubMed Central PMCID: 3853586.

46. Szyf M. Nongenetic inheritance and transgenerational epigenetics. Trends in molecular medicine. 2015 Feb;21(2):134–44. PubMed PMID: 25601643.

47. Vassoler FM, Sadri-Vakili G. Mechanisms of transgenerational inheritance of addictive-like behaviors. Neuroscience. 2014 Apr 4;264:198–206. PubMed PMID: 23920159. PubMed Central PMCID: 3872494.

48. Young LE, Fairburn HR. Improving the safety of embryo technologies: possible role of genomic imprinting. Theriogenology. 2000 Jan 15;53(2):627–48. PubMed PMID: 10735055.

49. Schneider E, Pliushch G, El Hajj N, Galetzka D, Puhl A, Schorsch M, et al. Spatial, temporal and interindividual epigenetic variation of functionally important DNA methylation patterns. Nucleic acids research. 2010 Jul;38(12):3880–90. PubMed PMID: 20194112. PubMed Central PMCID: 2896520.

50. Young LE. Imprinting of genes and the Barker hypothesis. Twin research: the official journal of the International Society for Twin Studies. 2001 Oct;4(5):307–17. PubMed PMID: 11869481.

51. Liu J, Hu X, Chen J, Li X, Wang L, Wang B, et al. Pericentral hepatocytes produce insulin-like growth factor-2 to promote liver regeneration during selected injuries in mice. Hepatology. 2017 Dec;66(6):2002–15. PubMed PMID: 28653763.

52. Song Y, Yang L. Transgenerational pancreatic impairment with Igf2/H19 epigenetic alteration induced by p,p’-DDE exposure in early life. Toxicology letters. 2017 Oct 5;280:222–31. PubMed PMID: 28867213.

53. Han L, Lee DH, Szabo PE. CTCF is the master organizer of domain-wide allele-specific chromatin at the H19/Igf2 imprinted region. Molecular and cellular biology. 2008 Feb;28(3):1124–35. PubMed PMID: WOS:000252606100023. English.

54. Engel N, Thorvaldsen JL, Bartolomei MS. CTCF binding sites promote transcription initiation and prevent DNA methylation on the maternal allele at the imprinted H19/Igf2 locus. Hum Mol Genet. 2006 Oct 1;15(19):2945–54. PubMed PMID: WOS:000240696100011. English.

55. Pathak S, Saxena M, D’Souza R, Balasinor NH. Disrupted imprinting status at the H19 differentially methylated region is associated with the resorbed embryo phenotype in rats. Reprod Fert Develop. 2010;22(6):939–48. PubMed PMID: WOS:000279384100006. English.

56. Zhang W, Yang J, Lv Y, Li S, Qiang M. Paternal benzo[a]pyrene exposure alters the sperm DNA methylation levels of imprinting genes in F0 generation mice and their unexposed F1-2 male offspring. Chemosphere. 2019 Apr 28;228:586–94. PubMed PMID: 31059956.

57. Bonham LW, Geier EG, Steele NZR, Holland D, Miller BL, Dale AM, et al. Insulin-Like Growth Factor Binding Protein 2 Is Associated With Biomarkers of Alzheimer’s Disease Pathology and Shows Differential Expression in Transgenic Mice. Front Neurosci-Switz. 2018 Jul 16;12. PubMed PMID: WOS:000438754300002. English.

58. Lewitt MS, Boyd GW. The Role of Insulin-Like Growth Factors and Insulin-Like Growth Factor-Binding Proteins in the Nervous System. Biochemistry insights. 2019;12:1178626419842176. PubMed PMID: 31024217. PubMed Central PMCID: 6472167.

59. Arai T, Busby W, Jr., Clemmons DR. Binding of insulin-like growth factor (IGF) I or II to IGF-binding protein-2 enables it to bind to heparin and extracellular matrix. Endocrinology. 1996 Nov;137(11):4571–5. PubMed PMID: 8895319.

60. Ly A, Ishiguro L, Kim D, Im D, Kim SE, Sohn KJ, et al. Maternal folic acid supplementation modulates DNA methylation and gene expression in the rat offspring in a gestation period-dependent and organ-specific manner. The Journal of nutritional biochemistry. 2016 Jul;33:103–10. PubMed PMID: 27152636.

61. El Hajj Chehadeh S, Dreumont N, Willekens J, Canabady-Rochelle L, Jeannesson E, Alberto JM, et al. Early methyl donor deficiency alters cAMP signaling pathway and neurosteroidogenesis in the cerebellum of female rat pups. American journal of physiology Endocrinology and metabolism. 2014 Dec 1;307(11):E1009–19. PubMed PMID: 25294213. PubMed Central PMCID: 4254983.

62. Pourie G, Martin N, Bossenmeyer-Pourie C, Akchiche N, Gueant-Rodriguez RM, Geoffroy A, et al. Folate-and vitamin B12-deficient diet during gestation and lactation alters cerebellar synapsin expression via impaired influence of estrogen nuclear receptor alpha. FASEB journal: official publication of the Federation of American Societies for Experimental Biology. 2015 Sep;29(9):3713–25. PubMed PMID: 26018677.

63. Daniel JM. Estrogens, estrogen receptors, and female cognitive aging: the impact of timing. Hormones and behavior. 2013 Feb;63(2):231–7. PubMed PMID: 22587940.

64. Saravia FE, Beauquis J, Revsin Y, Homo-Delarche F, de Kloet ER, De Nicola AF. Hippocampal neuropathology of diabetes mellitus is relieved by estrogen treatment. Cellular and molecular neurobiology. 2006 Jul-Aug;26(4-6):943–57. PubMed PMID: 16807785.

65. Cui J, Shen Y, Li R. Estrogen synthesis and signaling pathways during aging: from periphery to brain. Trends in molecular medicine. 2013 Mar;19(3):197–209. PubMed PMID: 23348042. PubMed Central PMCID: 3595330.

66. McEwen B, Akama K, Alves S, Brake WG, Bulloch K, Lee S, et al. Tracking the estrogen receptor in neurons: implications for estrogen-induced synapse formation. Proceedings of the National Academy of Sciences of the United States of America. 2001 Jun 19;98(13):7093–100. PubMed PMID: 11416193. PubMed Central PMCID: 34628.

